# Scop3P-Toolkit: executable structure-aware workflows linking PTMs, peptides, and mutations to protein function

**DOI:** 10.64898/2026.08.04.742789

**Authors:** Adrián Díaz, Natalia Tichshenko, Boris Depoortere, Rafael Andrade Buono, Paul De Geest, Wim F. Vranken, Lennart Martens, Pathmanaban Ramasamy

## Abstract

Post-translational modifications (PTMs) and genetic variants regulate protein function, signalling, and disease, but their interpretation requires integration of sequence annotations with structural, interaction, and biophysical context. Although resources such as Scop3P, UniProt, the Protein Data Bank, and AlphaFold provide extensive annotations and structural information, integrating these data into reproducible structure-aware analyses still requires custom scripting and manual coordination between multiple independent tools. To address this challenge, we developed **Scop3P-Toolkit**, an open-source executable analytical environment for interactive analysis of PTMs, mutations, and proteomics-derived peptides in their structural context. The toolkit integrates protein annotation retrieval with structural mapping, residue interaction network analysis, comparative structural analysis, and residue-level biophysical profiling within a unified framework. Experimentally supported phosphosites, phosphopeptides, and phosphoproteomics evidence are provided for human proteins through Scop3P, with optional integration of curated UniProt PTM annotations. UniProt-derived PTMs, sequence features, and genetic variants are available for proteins from any species, extending the framework beyond the human phosphoproteome. Scop3P-Toolkit supports structure-centric analyses including interpretation of PTMs and disease-associated variants, analysis of residue interaction networks and their rewiring across alternative conformations, structural localisation of peptides, and exploration of protein–protein, protein–ligand, and host–pathogen interfaces. Interactive visualisation links sequence annotations, three-dimensional structures, residue interaction networks, and biophysical profiles, enabling coordinated exploration across multiple molecular representations. The toolkit is distributed as Jupyter notebooks, browser-based Voilà applications, and a Galaxy interactive tool, providing transparent, accessible, and reproducible workflows for both computational and experimental researchers. By integrating biological annotation resources into executable, structure-aware workflows, Scop3P-Toolkit enables reproducible interpretation of PTMs, mutations, and proteomics data.

## Introduction

Post-translational modifications (PTMs) and genetic variants are major determinants of protein function. Phosphorylation, acetylation, ubiquitination, and related modifications regulate protein activity, molecular recognition, stability, localisation, and signalling^1^, whereas genetic variants can influence many of the same processes by altering residue chemistry, protein folding, interaction interfaces, or regulatory sites^2^. Although PTMs and variants are typically catalogued at the sequence level, their biological effects depend on three-dimensional structure, conformational state, solvent accessibility, residue interaction networks, and local biophysical properties^3–5^. Interpreting the functional consequences of a modification or variant therefore requires placing it within its structural context.

Although the information required to interpret PTMs and variants in their structural context is now available, it remains fragmented across multiple independent resources. UniProt provides curated protein sequences together with functional annotations across species^6^. The Protein Data Bank (PDB) is the primary archive of experimentally determined structures^7^, while AlphaFold and the AlphaFold Protein Structure Database have greatly expanded structural coverage across the proteome^8,9^. PDBe-KB^10^ further integrates structural and functional annotations onto these structures. For human phosphoproteins, Scop3P^11^ provides experimentally supported phosphosites, phosphopeptides, and associated phosphoproteomics evidence within structural and biophysical context, whereas UniProt supplies PTM and disease-variant annotations through its REST API services. Individually these resources are comprehensive, but they are primarily designed for data retrieval and visualisation rather than integrated analyses. Mapping PTMs onto structures, comparing them with disease variants, exploring residue neighbourhoods, constructing residue interaction networks, analysing biophysical profiles, and generating reproducible output still requires users to combine multiple resources with custom scripts and manual data transfer. This fragmentation limits usability and presents a particular challenge for education and training, where transparent and reusable workflows are essential.

Scop3P-Toolkit was developed to address this limitation by providing an executable environment for structure-centric analysis that integrates PTM, peptide and mutation annotations into reproducible workflows. For human phosphoproteins, the toolkit uses experimentally supported phosphosites, phosphopeptides, and phosphoproteomics evidence from Scop3P, with optional retrieval of curated UniProt PTM annotations. For proteins from any species, PTM, feature, and variant annotations are obtained from UniProt, extending the same analyses beyond the human phosphoproteome. Scop3P-Toolkit is available as a collection of Jupyter Notebooks^12^, standalone Voilà web applications, and a Galaxy interactive tool^13^, supporting reproducible analyses, interactive exploration, and community training^14^. Together, they provide a unified framework for structure-centric interpretation of PTMs, mutations, and proteomics data.

## Results

### The Scop3P-Toolkit framework

Scop3P-Toolkit consists of three complementary layers: a data layer, an analytical layer and a deployment layer (Fig. 1). The data layer integrates experimentally supported human phosphosites, phosphopeptides, and phosphoproteomics evidence from Scop3P; PTM, feature, and variant annotations from UniProt; experimental structures and structural annotations from PDBe-KB; and predicted structures from AlphaFold. It also accepts user-supplied structures and peptide sequences. The analytical layer combines these inputs to generate residue-resolved outputs, including mapped PTMs, peptides, and mutations, together with structural neighbourhoods, residue interaction networks (RINs), structural alignments, and biophysical profiles. The deployment layer exposes the same workflows through Jupyter Notebooks, standalone Voilà web applications, and a Galaxy interactive tool.

**Figure 1.**
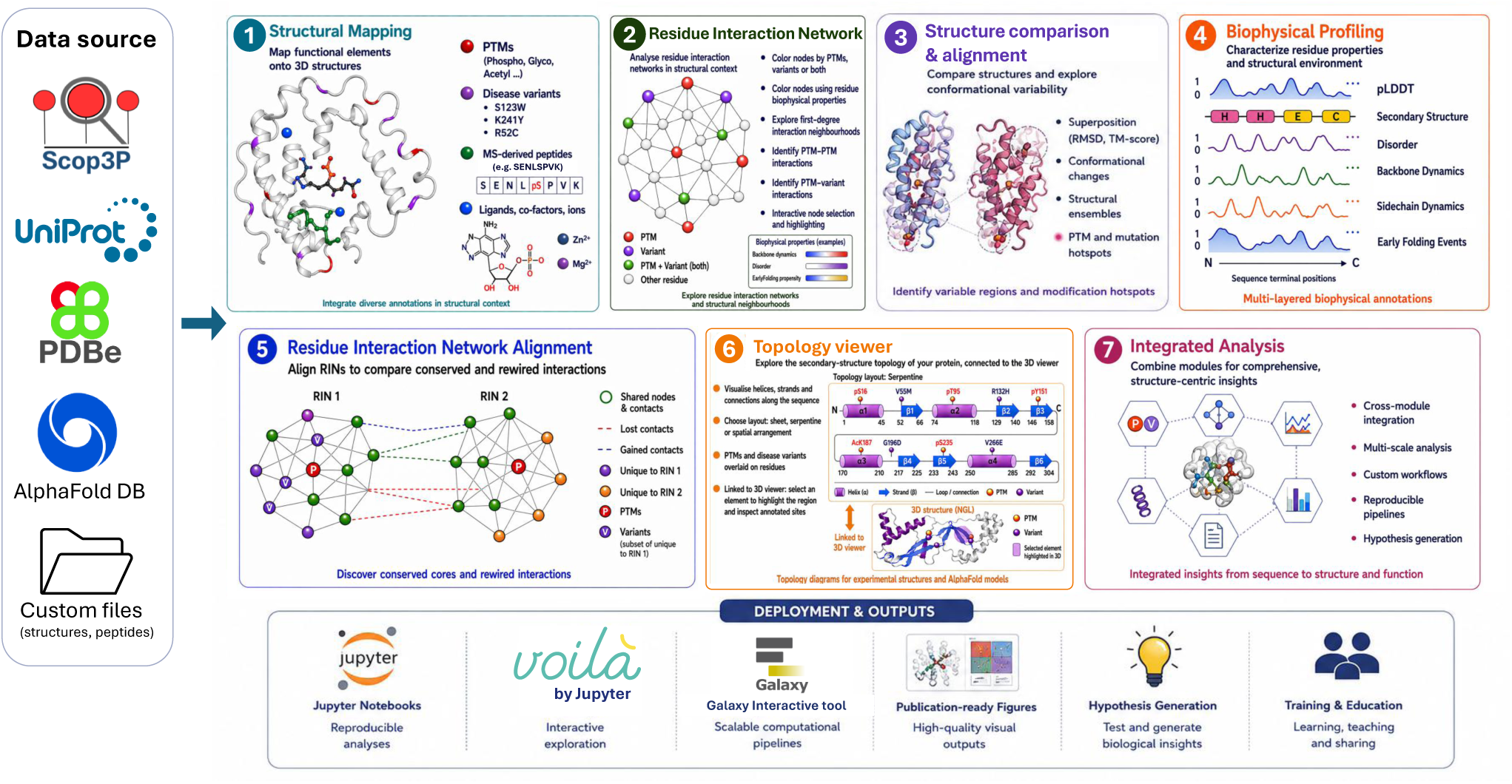
Conceptual overview of the Scop3P-Toolkit framework. Annotations are drawn from a set of data sources, including Scop3P, UniProt, the PDBe, the AlphaFold Database, and user-supplied structure or peptide files, and passed to seven analytical modules. (1) Structural mapping places PTMs, disease variants, mass-spectrometry-derived peptides, and ligands or cofactors onto three-dimensional structures. (2) Residue interaction network analysis explores networks in structural context, colouring nodes by PTMs, variants, or biophysical properties, resolving first-degree interaction neighbourhoods and PTM–PTM and PTM–variant contacts, with interactive node selection and highlighting. (3) Structure comparison and alignment superposes structures and explores conformational variability, covering conformational changes, structural ensembles, and PTM or mutation hotspots. (4) Biophysical profiling characterises residue properties and structural environment, including pLDDT, secondary structure, disorder, backbone and sidechain dynamics, and early-folding events. (5) Residue interaction network alignment compares two networks to identify conserved and rewired contacts, with PTMs and variants marked. (6) The topology viewer shows the secondary-structure topology of a protein, with sheet, serpentine, or spatial layouts, PTM and variant overlays, and linking to the three-dimensional viewer for both experimental and AlphaFold structures. (7) Integrated, cross-module analysis for reproducible, structure-centric insights from sequence to structure and function. The same workflows are deployed and shared as Jupyter notebooks, Voilà applications, and a Galaxy interactive tool, producing publication-ready figures and supporting hypothesis generation, training, and education.

Scop3P-Toolkit complements Scop3P by transforming curated phosphoproteomics annotations into executable, structure-centric workflows. Scop3P provides reprocessed phosphoproteomics evidence for human phosphosites, while UniProt provides PTM, feature, and variant annotations across species. These resources feed workflows for structural mapping, structural comparison, residue interaction network analysis, and interactive visualisation. For proteins beyond the human phosphoproteome, the same workflows operate on UniProt-derived annotations while retaining the enhanced support provided by Scop3P for human phosphoproteins. To handle incomplete datasets, the structure workflow first reports whether PTMs, variants, an AlphaFold model, and experimental structures are available for a given protein. Automatic retrieval is performed only when the corresponding resources are available, whereas manual PDB entry and structure upload remain possible in all cases.

The toolkit comprises five interactive workflows covering structure and PTM visualisation, peptide mapping, mutation-effect analysis, residue interaction network alignment, and secondary-structure topology (Table 1). Each workflow is available as a Jupyter Notebook and a standalone Voilà application. All workflows are bundled in the `bio2byte/scop3p-toolkit` Docker image, while the Galaxy interactive tool provides a single entry point for launching and switching between workflows.

**Table 1.** Interactive workflows available in Scop3P-Toolkit.

| Workflow | Purpose | Representative outputs |
| --- | --- | --- |
| Structure Visualisation | Interactive exploration of PTMs, peptides, variants, protein structures, residue interaction networks, and structural comparisons | Interactive structures, network views, and structural alignments |
| Peptide Mapper | Map proteomics-derived peptides and modified residues onto experimental or predicted protein structures | Annotated structures and peptide-mapping exports |
| Mutation Effect | Compare wild-type and mutant proteins using residue-level biophysical predictions | Comparative biophysical profiles and residue-level interpretation |
| RIN Alignment | Compare residue interaction networks between alternative structures or different proteins | Aligned networks, contact maps and similarity scores |
| Topology Viewer | Visualise two-dimensional secondary-structure topology linked to interactive three-dimensional structures | 2D topology with annotations, coupled to the 3D structure |

### From annotations to structural context

Every analysis in the toolkit begins from a single protein identifier. Given a UniProt accession, the toolkit retrieves the canonical protein sequence together with PTM, feature, and variant annotations, identifies available experimental and predicted structural models, and normalises all information to a common residue coordinate system. Human phosphosites, phosphopeptides, and associated phosphoproteomics evidence are retrieved from Scop3P through its REST API, while UniProt provides PTM, feature and variant annotations across species. Experimental structures from PDBe-KB and predicted structures from AlphaFold are mapped onto the canonical sequence, allowing modifications, variants, and peptides to be placed in their structural context (Fig. 2).

**Figure 2.**
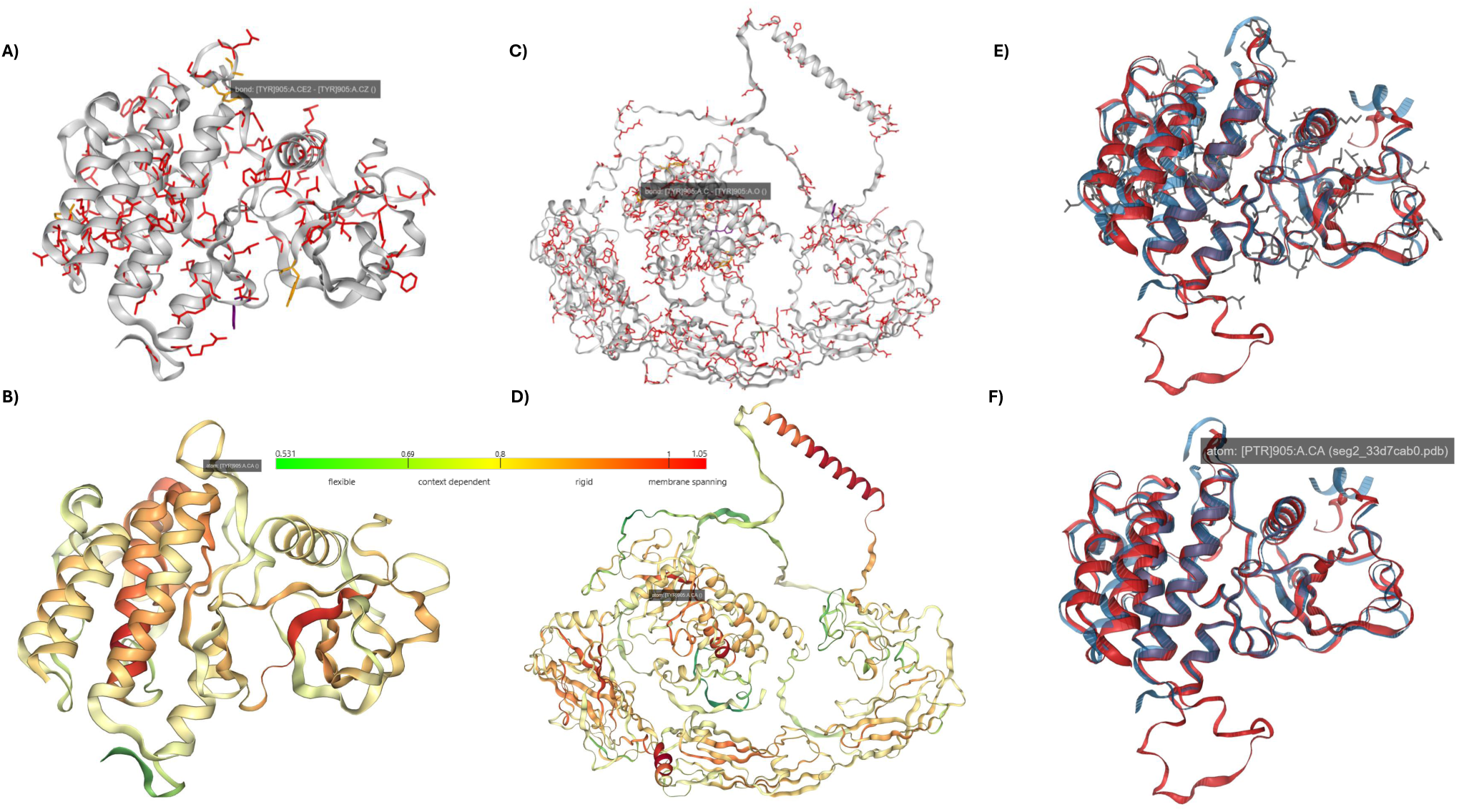
Structure and biophysical visualisation of RET (UniProt P07949) in Scop3P-Toolkit. (A, C) Post-translational modifications (yellow), disease mutations (red), and residues that are both modified and mutated (green; not shown in these panels) mapped onto (A) the experimental structure PDB 2IVS (chain A) and (C) the full-length AlphaFold model; the activation-loop residue Tyr905 is indicated. (B, D) The same two structures coloured by Bio2Byte backbone-dynamics values, using the scale shown from flexible (green, *∼*0.53) through context-dependent and rigid to membrane-spanning (red, *∼*1.05), so the mapped sites can be read in their biophysical context. (E, F) TM-align superposition of PDB 2IVS (chain A) and the AlphaFold model (shown in blue and red, respectively), with the phosphorylated activation-loop tyrosine (phospho-Tyr905) indicated in (E) and the mapped PTM and variant sites highlighted as grey sticks in (F).

This step comes first because every downstream analysis depends on it. Most interpretation questions begin with a sequence coordinate but can only be answered in three dimensions: a phosphosite, mutation, or peptide position must first be linked to the corresponding residue in a structure before accessibility, neighbouring residues, interaction networks, or conformational variability can be examined. Once the mapping is established, all subsequent analyses proceed from the same identifier, apart from the user’s choice of structure. Residue-level biophysical properties are predicted directly from the canonical sequence, residue interaction networks are generated from the selected AlphaFold model or experimental structure, and structural comparison, peptide mapping, and mutation analysis are performed on the same residue axis, eliminating manual file handling and identifier conversion.

Residue mapping is therefore treated as an explicit, verifiable step rather than an implicit component of structural visualisation. The resulting mapping tables and residue coordinates remain accessible throughout the analysis, allowing them to be checked, reused and exported for downstream applications.

### Structure-guided analysis of PTMs and variants

Once structural context has been established, PTMs and variants can be examined within a common structural and biophysical framework. For human proteins, experimentally observed phosphosites and their associated phosphoproteomics evidence are retrieved from Scop3P^11^ and combined with PTM, feature, and variant annotations from UniProt, while for proteins from other species the same analyses operate on UniProt annotations alone. Either an experimental structure from PDBe-KB or a predicted AlphaFold model can be selected, allowing PTMs and disease variants to be explored across experimental and predicted structures. Rather than presenting static annotation tracks, the toolkit places each residue within its structural environment, where solvent accessibility, local residue neighbourhoods, nearby variants, residue-level biophysical properties, and differences between alternative conformations can be examined together.

Residue-level biophysical properties are predicted using the sequence-based predictors implemented in the b2bTools package^5,15^, DynaMine (backbone dynamics, side-chain dynamics, and helix, sheet, and coil propensities), DisoMine (disorder propensity), and EFoldMine (early-folding propensity). A structure can be coloured by backbone dynamics, side-chain dynamics, disorder propensity, early-folding propensity, or predicted helix, sheet, or coil propensity, while PTM and mutation sites are highlighted, allowing predicted residue behaviour to be interpreted in its structural context. The toolkit does not infer function automatically; instead, it provides the structural and experimental evidence needed to formulate and evaluate hypotheses about the functional relevance of individual sites (Fig. 2).

Structural comparison is integrated within the same workflow. TM-align^16^ is used to superpose two structures, such as an experimental structure and its corresponding AlphaFold model or two alternative experimental conformations, reporting the root-mean-square deviation (RMSD) and template-modelling score (TM-score) of the alignment. PTMs and variants can be overlaid on either or both structures, with the view optionally restricted to the aligned region (Fig. 2, panels E and F). This enables conformational differences to be assessed alongside PTM and mutation distributions, helping users determine whether structural rearrangements coincide with modified or variant-rich regions. A complementary comparison at the network level, reporting residue contacts that are conserved, lost or gained between the same structures, is provided by the residue interaction network alignment workflow described below (Fig. 6).

### Structural localisation of mass spectrometry-derived peptides

A dedicated workflow maps peptides onto the same residue coordinate system, projecting phosphopeptides from Scop3P or user-supplied peptide datasets onto experimental or predicted protein structures. Peptide coverage can be inspected along the protein sequence and visualised in three dimensions, enabling residue accessibility and structural localisation to be assessed (Fig. 3). Modified residues identified within peptides are highlighted automatically, and when multiple peptides are selected, shared and unique sequence regions are identified to facilitate comparison. Mapping peptides onto protein structures supports the evaluation of signature peptides for targeted proteomics and diagnostics, and of other functional peptides such as immunopeptides, where structural accessibility and the surrounding molecular environment can influence biological function.

**Figure 3.**
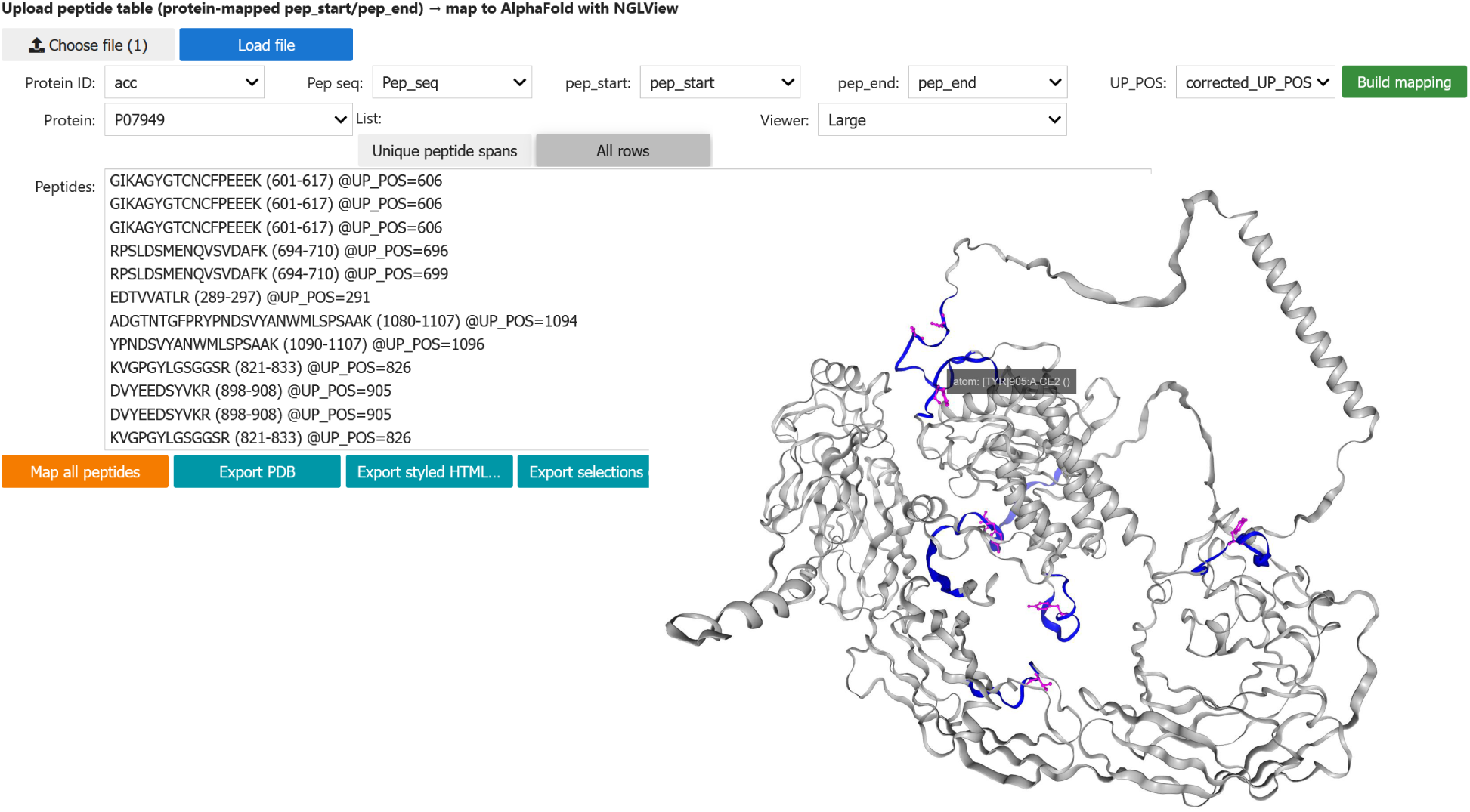
Peptide Mapper workflow of Scop3P-Toolkit, shown for RET (UniProt P07949). Peptides obtained from Scop3P or uploaded by the user as a table are mapped onto the AlphaFold structure with NGLView; the selected peptides are highlighted in blue and the phosphosites they contain are shown as magenta sticks (the tooltip indicates Tyr905, within the mapped peptide DVYEEDSYVKR, residues 898–908). The peptide list, the choice of unique spans or all rows, and export of the mapped structure as PDB or as a styled standalone HTML session are available from the controls on the left.

### Mutation-effect analysis

The mutation workflow compares wild-type and user-defined mutant proteins on the same residue coordinate system. One or more amino acid substitutions can be introduced interactively to generate a mutant sequence, after which all residue-level biophysical properties are recalculated with the b2bTools predictors. Wild-type and mutant profiles are displayed as interactive residue-wise profiles together with an inference step summarising the predicted changes in biophysical behaviour, and where structural models are available, these changes can be interpreted directly in their structural context. Because PTMs, mutations, and structural features share the same coordinate system, substitutions can be evaluated in relation to nearby modification sites, interaction interfaces and predicted structural properties. This enables users to examine, for example, whether a mutation adjacent to a phosphosite alters local dynamics or accessibility, or whether substitutions within structured regions influence predicted folding behaviour or flexibility (Fig. 4).

**Figure 4.**
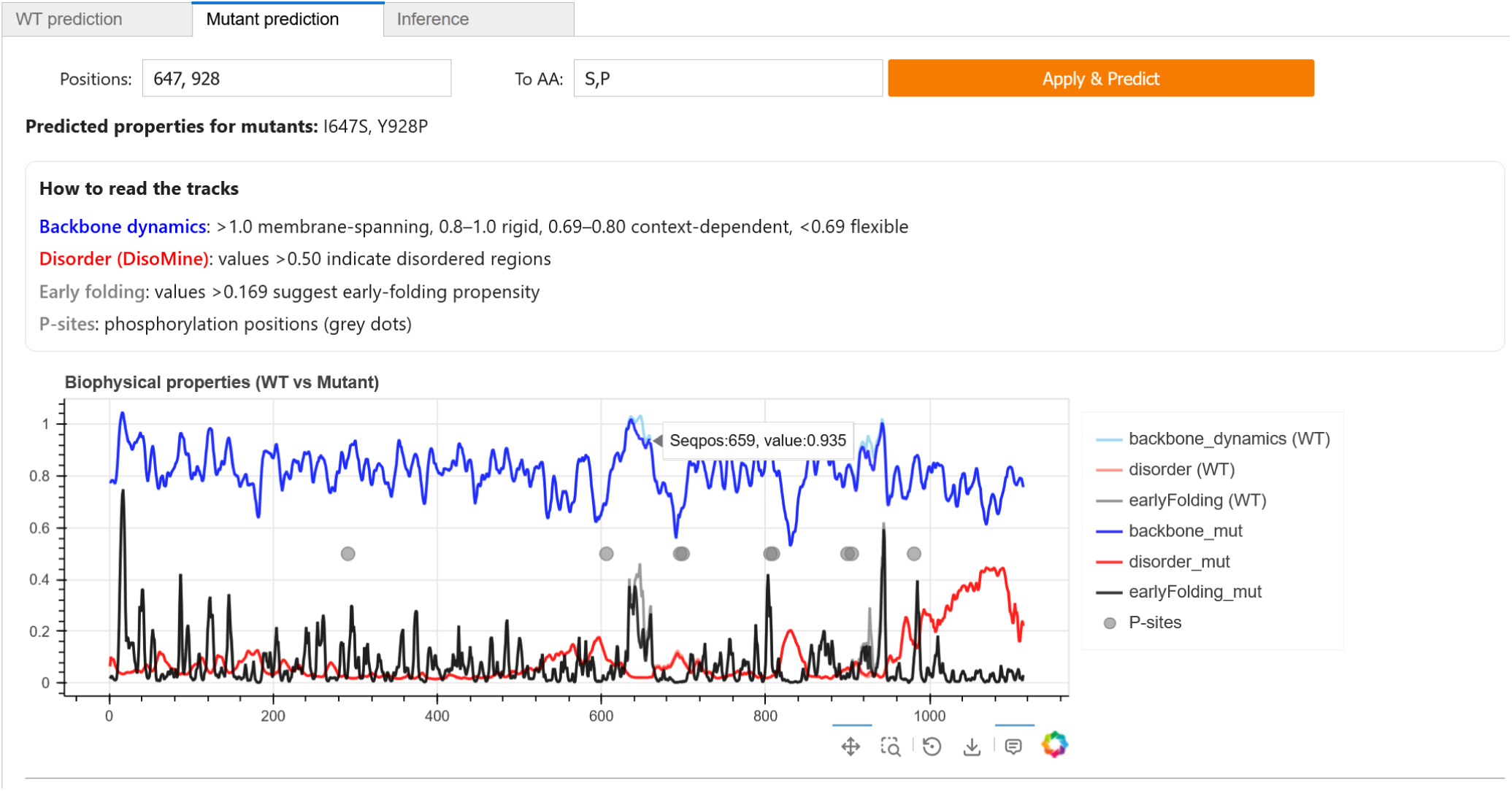
Mutation Effect workflow of Scop3P-Toolkit, shown for RET (UniProt P07949). Residue-level biophysical properties predicted with b2btools, backbone dynamics (blue), disorder (red), and early folding (black), are compared between the wild-type protein and a user-defined mutant along the full sequence; here the substitutions I647S and Y928P are applied. Phosphorylation sites are marked as grey dots, and the guide lists the interpretation thresholds for each property. The Inference tab (not shown) summarises the wild-type-to-mutant change at each mutated position and within a *±*5 residue window, reporting value shifts and any change of biophysical class (for example membrane-spanning to rigid, or early-folding to non-early-folding).

### Residue interaction networks and their alignment

Three-dimensional protein structures can be represented as residue interaction networks (RINs), where residues form the nodes and spatial contacts define the edges. Scop3P-Toolkit generates RINs from selected AlphaFold models, experimental structures retrieved through PDBe-KB, or user-supplied structures, allowing the same protein to be explored as a network across different structural contexts. PTM and mutation sites are highlighted within the network and their local interaction neighbourhoods can be inspected directly. The same residue-level biophysical properties used for structure colouring can also be projected onto network nodes, while node borders and sizes indicate PTM and variant status. This integrates residue connectivity, predicted biophysical behaviour, and molecular annotations within a single representation (Fig. 5). Compared with direct three-dimensional visualisation, RINs emphasise residue connectivity and facilitate exploration of local connectivity, allowing interpretation to extend beyond residue identity to its network context. Network construction uses established Python libraries, including Biopython for structure parsing and NetworkX, Pandas, and NumPy for graph analysis^17–20^.

**Figure 5.**
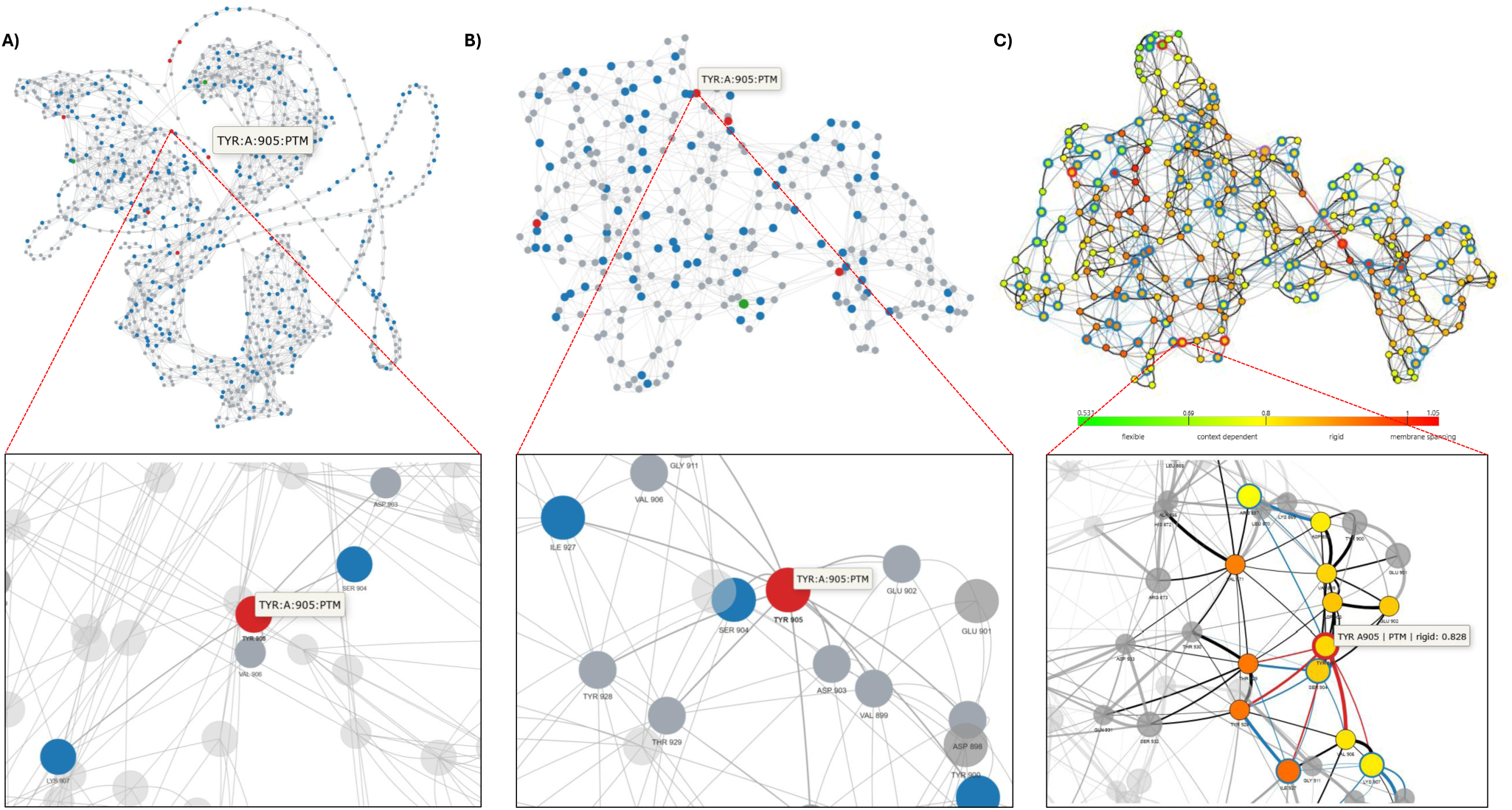
Residue interaction network (RIN) analysis of RET (UniProt P07949) in Scop3P-Toolkit. (A, B) RINs built from (A) the AlphaFold model and (B) the experimental structure PDB 2IVS (chain A), with nodes coloured by annotation status: PTMs in red, mutations in blue, and residues that are both in green; unannotated residues are grey. (C) The RIN with node fill coloured by Bio2Byte backbone-dynamics values, using the scale shown from flexible (green) through context-dependent and rigid to membrane-spanning (red), while node borders keep the PTM, mutation, and both colouring of (A) and (B), so that topology, biophysical property, and annotation appear in a single view. Bottom row: for each network, a zoom on the first-degree contacts of the phosphorylated activation-loop tyrosine Tyr905, with neighbouring residues labelled; in (C) the tooltip reports residue status and backbone class and value (Tyr905, PTM, rigid, 0.828).

**Figure 6.**
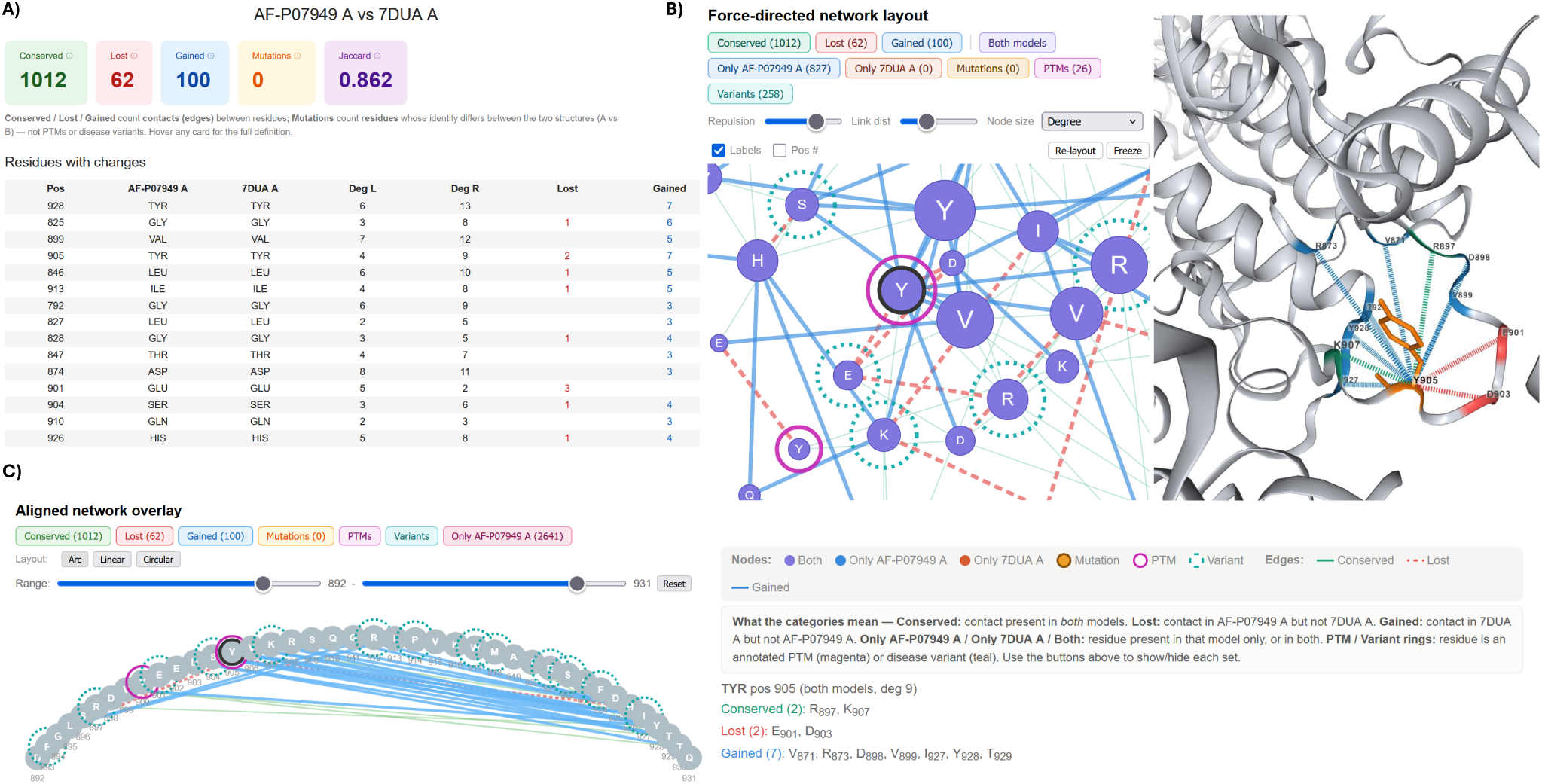
Residue interaction network alignment in Scop3P-Toolkit, comparing the AlphaFold model of RET (UniProt P07949, chain A) with the experimental structure PDB 7DUA (chain A). (A) Summary of the comparison: counts of conserved, lost, and gained contacts and mutated positions, a Jaccard network-similarity score (0.862), and a table of residues whose contacts change, listing the per-structure node degree and the number of lost and gained contacts. (B) Force-directed network layout linked to a three-dimensional structure view. Nodes are coloured by presence in both structures or in one only, with rings marking annotated PTMs (magenta) and disease variants (teal); edges are coloured as conserved (green), lost (red dashed), or gained (blue), and category buttons show or hide each set. Selecting a node highlights the corresponding residue and its contacts on the structure (right), here the phosphorylated activation-loop tyrosine Tyr905 with its conserved, lost and gained contacts. (C) Aligned network overlay with arc, linear, or circular layouts and a sequence-range slider, showing the contact changes within a selected region.

Scop3P-Toolkit also provides a dedicated workflow for comparing residue interaction networks. For a selected protein, networks generated from an AlphaFold model and available experimental structures can be compared using a user-defined contact distance cutoff. Two comparison modes are supported. In the same-protein mode, residues are matched by sequence position to identify conserved, lost, and gained residue contacts together with positions that differ in residue identity. In the cross-protein mode, networks are aligned using Weisfeiler–Lehman neighbourhood signatures^21^, residue-type similarity, and node degree, enabling different proteins to be compared on the basis of their network topology.

The comparison is summarised by the numbers of conserved, lost, and gained contacts, mutated positions, and a Jaccard network similarity score, while residues involved in rewired contacts are reported together with their network properties. Complementary interactive network views, such as a force-directed layout and an aligned overlay, together with contact maps, allow connectivity changes to be explored at both global and residue level, and a linked three-dimensional view highlights the corresponding residues and interaction partners directly on the structure. PTM and variant annotations are retained throughout the comparison, enabling structural rewiring associated with conformational changes or amino acid substitutions to be interpreted at the level of individual residue contacts (Fig. 6).

### Secondary-structure topology

A complementary two-dimensional topology view is generated from the structure’s secondary-structure records, or DSSP^22^ assignments when these are absent, relating helices, strands, and their connections to features along the protein sequence (Fig. 7). Elements can be arranged in sheet, serpentine, or spatial-arrangement layouts, and PTM and disease-variant annotations are overlaid on the corresponding residues, with a site-probability filter for Scop3P evidence. Topology diagrams are available for both experimental structures and predicted AlphaFold models. Each diagram is linked directly to the NGL three-dimensional viewer, so that selecting a topology element highlights the corresponding region on the structure and lets its annotated sites be inspected together. The current view can be exported as JSON for reproducibility. Extending topology diagrams to predicted structures, which existing visualisation tools rarely support, lets secondary-structure context be read alongside the same PTMs, variants, and structural views used throughout the toolkit, keeping every representation anchored to one residue coordinate system. Conventional topology viewers are oriented towards deposited PDB entries, and neither the PDBe^10^ nor the AlphaFold Database^8,9^ provides an equivalent 2D topology view for its models, leaving this representation largely unavailable for predicted structures.

**Figure 7.**
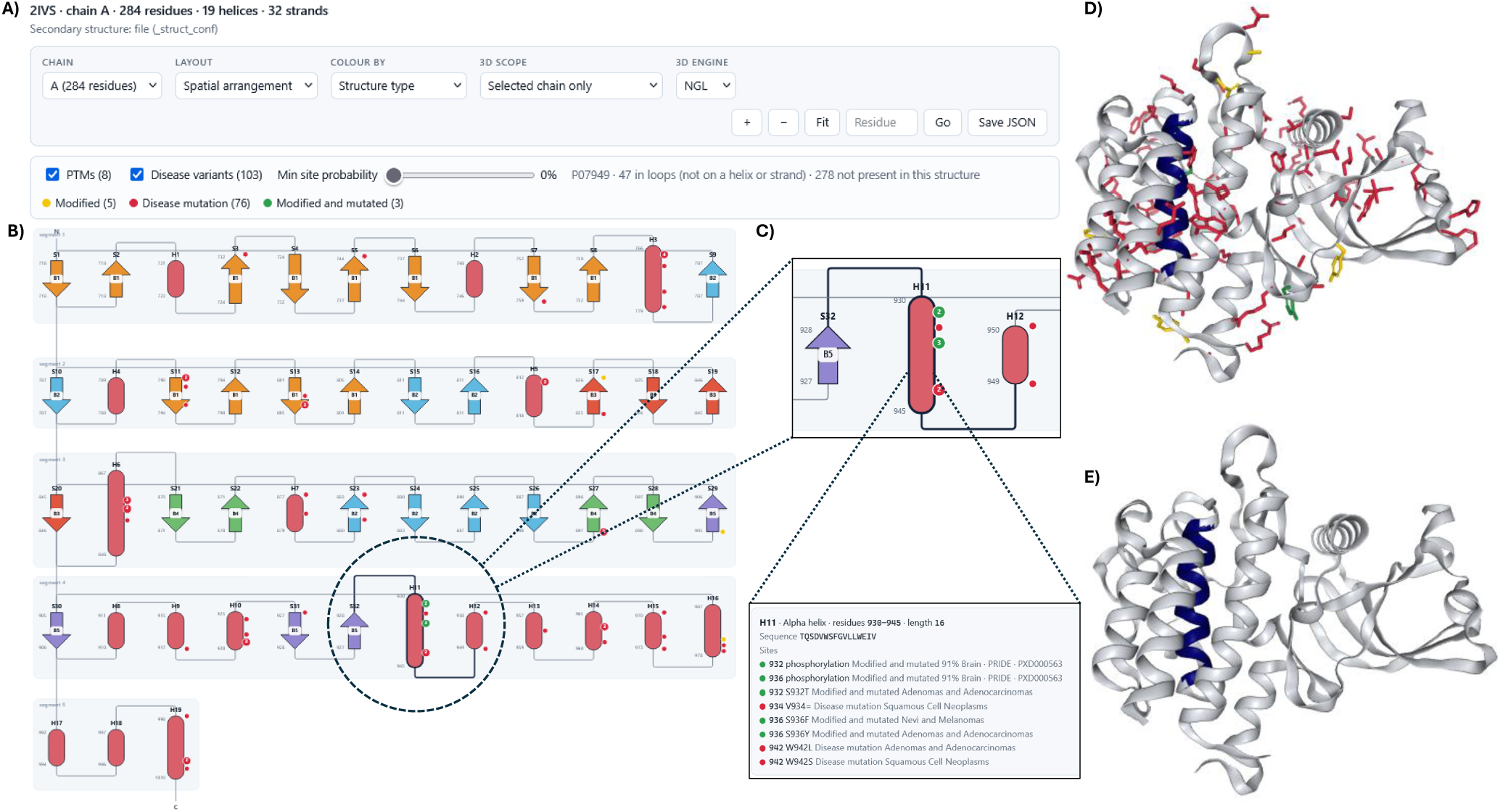
Secondary-structure topology viewer of Scop3P-Toolkit, shown for RET (UniProt P07949), experimental structure PDB 2IVS (chain A). Secondary structure is read from the structure’s own records where available and computed with DSSP otherwise. (A) Control bar and annotation panel: the control bar sets the chain, the two-dimensional layout (sheet, serpentine, or spatial arrangement), the colouring, and the 3D view, and exports the current view as JSON; PTM and disease-variant overlays can be toggled, a slider filters sites by minimum Scop3P site probability, and the legend distinguishes modified (yellow), disease-mutation (red), and modified-and-mutated (green) residues. (B) The two-dimensional topology diagram in the spatial-arrangement layout: *α*-helices as rounded bars and *β*-strands as arrows coloured by sheet membership, connected by loops with the N-and C-termini marked, and per-residue PTM and variant sites shown as coloured dots on the corresponding elements. (C) Two linked insets for a selected element, helix H11 (residues 930–945): a zoom of the element with its PTM and variant sites marked as coloured dots, and a detail panel listing each site with its residue, modification or variant type, and supporting evidence (here phosphorylation at 932 and 936, and disease mutations including W942L and W942S). (D, E) The same chain rendered as a three-dimensional structure (NGL), with the mapped sites shown as sticks, modified (yellow), disease-mutation (red), and modified-and-mutated (green), and the element selected in (C), helix H11, highlighted in blue; shown with (D) and without (E) the site overlay.

### Applications across the structure–function axis

The strength of Scop3P-Toolkit lies in bringing experimental proteomics evidence, structural information, residue interaction networks and residue-level biophysical properties into a common structural framework, allowing multiple sources of evidence to be interpreted together rather than in isolation. We illustrate this using RET (UniProt P07949), a receptor tyrosine kinase whose activity is regulated by phosphorylation of its activation loop and whose germline and somatic mutations are associated with multiple endocrine neoplasia type 2 and thyroid cancer^23^. RET represents a biologically relevant example in which phosphorylation, genetic variation and conformational change converge on the same protein, making it an ideal system for demonstrating the integrated analyses provided by Scop3P-Toolkit.

Starting from a UniProt accession, the toolkit retrieves RET phosphosites and maps activation-loop and C-terminal tyrosines onto both experimental and AlphaFold structures (Fig. 2), while the phosphopeptides carrying these sites are localised on the corresponding structures (Fig. 3). Phosphorylation of the activation-loop tyrosines is associated with stabilisation of the active kinase conformation^24^, making the structural behaviour of these sites across different conformations of particular interest. Residue-level biophysical properties are projected onto both the three-dimensional structure and the corresponding residue interaction network (Fig. 5), while network alignment identifies residue contacts that are conserved, lost, or gained between conformations (Fig. 6). User-defined substitutions at phosphorylatable and neighbouring residues, such as I647S and Y928P, are compared with the wild-type protein (Fig. 4), and clinically annotated RET variants are overlaid within the same residue coordinate system. Together, these analyses provide a structure-centric interpretation of how phosphorylation, genetic variation, and conformational change relate to RET function.

RET also illustrates why analysing a single structure is often insufficient. The activation loop, which contains the phosphotyrosine cluster centred on Tyr905 together with neighbouring sites at Tyr909 and Tyr928, adopts multiple conformations that depend on its phosphorylation state. Previous work showed that available RET structures separate into distinct conformational states, including a phosphorylation-associated open activation-loop conformation that is not consistently reproduced by AlphaFold models, including the phosphorylation-aware AF3-p variant^25^. Interpreting these phosphosites therefore requires comparison across multiple conformations rather than reliance on a single structural model. Scop3P-Toolkit is designed to support this type of analysis by maintaining the same residue coordinate system throughout all workflows. The activation-loop residues remain linked across structural visualisation, biophysical profiling, residue interaction networks, network alignment, and mutation analysis (Figs. 2, 5 and 6), allowing users to move directly between complementary structural representations while following the same biologically relevant sites. Rather than committing to a single structural model, each available conformation can be examined individually, compared directly, and interpreted within its own structural and interaction context.

Although illustrated here using RET, the same environment supports a broad range of applications beyond phosphosite annotation. Mapping peptides onto protein structures reveals whether they are surface-exposed and defines their local structural environment, supporting the evaluation of signature peptides for targeted proteomics, diagnostic applications, and other functional peptides such as immunopeptides. Mapping PTMs and variants relative to ligand-binding regions places modifications and mutations into a drug-binding context, while examining PTM and mutation hotspots at protein–protein or host– pathogen interfaces helps identify sites that may influence molecular recognition. In each case, the same integrated environment is reused to address different biological questions, allowing experimental proteomics evidence, structural information, and residue-level biophysical properties to be interpreted together within a single, reproducible analytical framework.

### Deployment, reproducibility and training

A central aim of Scop3P-Toolkit is to make structure-centric analyses accessible without compromising reproducibility. Each workflow is available as a Jupyter notebook that exposes the complete analysis code for transparency and customisation^12^, and as a standalone Voilà application that provides the same functionality through a code-free browser interface. All workflows are bundled in the `bio2byte/scop3p-toolkit` Docker image and deployed as a Galaxy interactive tool^13^, making the toolkit available within an established, freely accessible platform without local installation, alongside a single entry point for launching and switching between workflows.

The toolkit is also used as a training resource. The associated ELIXIR-oriented training material guides users from protein sequence and structure to PTM and mutation interpretation through hands-on workflows, combining biological background with notebooks for biophysical prediction, PTM and peptide mapping, mutation-effect analysis, and structural comparison, while presenting Scop3P and the Bio2Byte tools as ELIXIR Belgium Node services^5,14^. The interactive visualisation components were further developed during BioHackathon Europe 2025, which prototyped a reusable library of linear, network, and structural views for PTM data^26^.

Providing each workflow as a notebook, a browser application and a Galaxy interactive tool lowers the barrier for experimental researchers while preserving complete analytical detail for computational users. At the same time, it supports FAIR practice by keeping data access, analysis steps, parameters, and outputs open, reproducible, and reusable. Results can also be shared outside the application through self-contained interactive HTML exports that preserve mapped PTMs, variants and structural colouring, allowing individual analyses to be saved, distributed, and reopened without rerunning the toolkit.

## Discussion

Scop3P-Toolkit addresses a practical gap in protein bioinformatics: the distance between biological annotation resources and executable analysis. Many databases provide PTM, mutation, and structural information, but interpreting these data typically requires users to integrate information from multiple resources through custom scripts or manual workflows. Scop3P-Toolkit provides a workflow-based analytical layer that connects these resources through reusable modules for structural mapping, PTM and peptide analysis, mutation-effect analysis, residue interaction networks, biophysical profiling, structural comparison, and visualisation. Scop3P-Toolkit is not a database. Scop3P remains the knowledge base that supplies experimentally supported human phosphosite information in context, while the toolkit uses these data within executable workflows and extends the same analytical framework to proteins from any species through UniProt-derived annotations.

Treating PTMs, peptides and mutations within a shared residue coordinate system is what makes the toolkit broadly applicable. The same analytical components that map a phosphosite can also localise a functional peptide on a protein surface, place a variant next to a ligand-binding region, or examine a modification at a host–pathogen interface. The RET example illustrates this principle by showing how phosphosites, mutations and conformational variability can be interpreted together rather than as separate annotations. More broadly, RET highlights a limitation of relying on any single structural model. Where a modification reshapes the conformational landscape, and where predicted structures preferentially represent the dominant state, meaningful interpretation requires comparison across multiple conformations rather than reliance on a single model. Enabling these comparisons across structural, network, and biophysical views within a shared residue coordinate system is where an executable toolkit provides clear advantages over inspecting individual structures. Across these applications, the toolkit does not assign fixed functional labels; instead, it assembles the structural and biophysical evidence needed to formulate and evaluate biological hypotheses, while preserving the complete analytical context in a reproducible form.

Several limitations should be considered. Structural interpretation depends on the availability and quality of experimental or predicted structures, and AlphaFold models, despite their broad coverage, may not capture all biologically relevant states, including PTM-regulated, ligand-bound, or highly flexible conformations. The toolkit partially mitigates missing structural information by reporting data availability and supporting manual structure upload, but meaningful structural interpretation ultimately depends on suitable models being available. Residue interaction networks represent simplified abstractions of molecular contacts and should therefore be interpreted alongside the underlying three-dimensional structures. Likewise, PTM annotations differ in their level of experimental support across resources and species, requiring appropriate consideration of the underlying evidence during biological interpretation.

In summary, Scop3P-Toolkit provides a reusable environment in which protein structures, residue-level biophysical properties, PTMs, peptides, and genetic variants can be analysed together. By integrating experimental proteomics evidence with structural and biophysical context through structured, shareable workflows, the toolkit extends the Scop3P ecosystem beyond database access while supporting reproducible research, community training, and FAIR data practices. As an open-source project, Scop3P-Toolkit is also intended to grow as a community platform, where users can request features, report needs, or contribute new analyses and functionality.

Looking ahead, Scop3P-Toolkit is designed to grow as a community project. Because every workflow is an open, self-contained notebook, users can adapt existing analyses, contribute new ones, or request features and changes through the public repository, and we welcome contributions from the wider structural-biology and proteomics communities. Natural directions for this shared development include broader PTM coverage, ensemble-aware structural interpretation, additional evidence layers such as evolutionary conservation and molecular interactions, deeper Galaxy integration, and on-demand structure prediction, so that proteins lacking an experimental or predicted model, including designed sequences, can be analysed within the same workflows.

## Methods

### Software organisation

Scop3P-Toolkit is implemented as a collection of Jupyter notebooks, standalone Voilà applications and a Galaxy interactive tool. The notebooks provide task-specific workflows for structure and PTM visualisation, peptide mapping, mutation-effect analysis, residue interaction networks and their alignment, and secondary-structure topology. All workflows are bundled in the `bio2byte/scop3p-toolkit` Docker image, enabling local execution or deployment through Galaxy.

### Annotation retrieval

For human phosphoproteins, phosphorylation annotations are retrieved from the Scop3P modifications REST API endpoint^11^. For proteins from any species, protein sequences, PTM and feature annotations, and disease variants are retrieved from UniProt through its REST API^6^. Retrieved annotations are normalised to residue positions in the canonical protein sequence to support structural mapping and visualisation.

### Structural data handling

Experimental structures and associated structural annotations are retrieved through PDBe-KB^10^, while predicted models are obtained from the AlphaFold Protein Structure Database^8,9^. Author residue numbering in experimental structures often differs from UniProt numbering, so the toolkit uses the PDBe updated mmCIF files and parses their SIFTS residue-level mappings to relate UniProt positions to PDB author residue numbers^10^. AlphaFold models follow UniProt residue numbering directly. Structural residues are thereby linked to canonical sequence coordinates so that PTMs, peptides, and mutations are displayed on the correct residues, while predicted confidence scores (e.g. pLDDT) are retained for interpretation where available.

### PTM and peptide mapping

PTM workflows project modified residues from sequence coordinates onto available experimental or predicted structures. Peptide workflows accept phosphopeptides retrieved from Scop3P or peptides uploaded by users. Selected peptide spans are mapped onto the structure, displaying combined peptide coverage together with shared and unique sequence regions and modified residues, allowing peptide coverage and modification sites to be interpreted alongside structural accessibility and local environment.

### Mutation-effect workflow

The mutation workflow generates altered protein sequences from one or more user-defined substitutions and compares wild-type and mutant profiles. Residue-level biophysical properties are recomputed for the mutant sequence with the b2bTools predictors (DynaMine, DisoMine, and EFoldMine) and visualised along the residue axis, allowing mutation positions to be interpreted together with PTM sites, peptides, structural features, and predicted properties. Wild-type and mutant profiles are displayed as interactive line plots rendered with the Bokeh Python library^27^, and an inference step reports whether a substitution changes the predicted biophysical state of a residue.

### Residue interaction network construction and alignment

Residue interaction networks are generated from AlphaFold models or PDB structures retrieved by the toolkit, or from user-uploaded structures. Structures are parsed with Biopython^17^; residues are represented as nodes and residue contacts as edges between residues whose C*α* atoms lie within a user-defined distance cutoff (8 Å by default), with C*β* atoms (C*α* for glycine) available as an alternative. Contacts between adjacent sequence neighbours are excluded. Networks are constructed with NetworkX^18^, while PTM and mutation sites are incorporated as node annotations. Nodes can be coloured according to selected residue-level biophysical properties, and node borders and sizes indicate PTM and variant status.

Two residue interaction networks can be compared directly. A position-based comparison matches residues by sequence position and reports maintained, lost, and gained contacts together with positions whose residue identity differs. A complementary topology-based alignment matches nodes using Weisfeiler–Lehman neighbourhood signatures^21^, residue-type similarity, and node degree, combines these measures into a node similarity matrix, and resolves the optimal assignment with SciPy^28^. Results are summarised as maintained, lost, and gained edge sets together with a Jaccard network-similarity score and visualised as contact maps, aligned networks, and force-directed layouts. A linked view pairs the force-directed network with an NGL structure viewer, allowing selected network nodes and their contacts to be highlighted directly on the corresponding three-dimensional structure.

### Biophysical prediction, structural comparison and topology

Residue-level biophysical properties are predicted from protein sequences using the sequence-based predictors implemented in the b2bTools package^5,15^, including DynaMine (backbone dynamics, side-chain dynamics, and helix, sheet, and coil propensities), DisoMine (disorder propensity), and EFoldMine (early-folding propensity). Predicted properties can be projected onto both three-dimensional structures and residue interaction networks. Structural comparison is performed with TM-align^16^, which superposes two structures and reports their RMSD and TM-score, supporting comparison of experimental and predicted models or alternative conformations, while PTM and variant annotations can be overlaid on either or both aligned structures, optionally restricted to the structurally aligned region. A two-dimensional topology view is generated for both experimental and predicted (AlphaFold) structures from their deposited secondary-structure records where available, falling back to DSSP^22^ assignments otherwise, and is linked to an NGL^29^ viewer, allowing selected topology elements to be highlighted directly on the corresponding three-dimensional structure.

Three-dimensional structures are rendered with NGL and py3Dmol^29,30^ and residue interaction networks with pyvis^31^, and the structural, network, and alignment views can be exported as self-contained HTML files that preserve the mapped annotations and visualisation settings.

## Data Availability

The Scop3P resource is publicly available at https://iomics.ugent.be/scop3p/. Protein sequences, PTM and feature annotations, disease variants, experimental structures and structural annotations, and predicted structures are retrieved from the public Scop3P, UniProt, PDBe-KB, and AlphaFold resources, respectively. Training material and example datasets are available through the associated GitHub repositories.

## Code Availability

Scop3P-Toolkit notebooks and applications are available at https://github.com/Bio2Byte/Scop3P-notebooks, released under the Apache License 2.0. The associated training material is available at https://github.com/Bio2Byte/training-protein-dynamics-post-translational-modifications, with rendered Voilà applications at https://bio2byte.github.io/training-protein-dynamics-post-translational-modifications/, under the CC BY 4.0 licence and archived at Zenodo (DOI: https://doi.org/10.5281/zenodo.18242776). The toolkit is deployed as a Galaxy interactive tool on both the European (https://usegalaxy.eu/?tool_id=interactive_tool_scop3p_toolkit) and Belgian (https://usegalaxy.be/?tool_id=interactive_tool_scop3p_toolkit) Galaxy servers.

## Acknowledgements

The authors thank all data submitters for making their data publicly available, and the developers of the external resources used in this study for providing open-access data and API platforms. P.R., A.D., and N.T. thank the participants of BioHackathon Europe 2025 for their contributions, and the authors thank the participants of the Scop3P training sessions for their valuable feedback.

## Author Contributions

P.R. conceived the toolkit and drafted the manuscript. P.R., A.D., and N.T. contributed to toolkit development, training material, and workflow design. A.D., B.D., R.A.B., and P.D.G. contributed to data infrastructure, software, and deployment. W.F.V., L.M., and P.R. supervised the project. All authors contributed to manuscript preparation and approved the final version.

